# TipN’s involvement with centromere segregation in Caulobacter crescentus

**DOI:** 10.1101/2023.12.20.572679

**Authors:** Morgan Letzkus, Corey Trela, Paola E. Mera

## Abstract

Bacteria’s ability to maintain chromosomal integrity throughout their life cycle is crucial for their survival. In *Caulobacter crescentus*, the polar factor TipN has been proposed to be involved with the partitioning system ParABS. However, cells with *tipN* knocked out display subtle *parS* segregation defects. We hypothesized that TipN’s role with *parS* segregation is obscured by other forces that are ParABS-independent. To test our hypothesis, we removed one of those forces – chromosome replication – and analyzed the role of TipN with ParA. We first demonstrate that ParA retains its ability to transport the centromeric region *parS* from the stalked pole to the opposite pole in the absence of chromosome replication. Our data revealed that in the absence of chromosome replication, TipN becomes essential for ParA’s ability to transport *parS*. Furthermore, we identify a potential connection between the replication initiator DnaA and TipN. Although TipN is not essential for viability, *tipN* knockout cells lose viability when the regulation of DnaA levels is altered. Our data suggest that the DnaA-dependent susceptibility of *tipN* knockout cells is connected to *parS* segregation. Collectively, this work provides insights into the complex regulation involved in the coordination of chromosome replication and segregation in bacteria.

## BACKGROUND

The ability of cells to successfully replicate and pass on their genetic information to their offspring is essential for all domains of life. Unlike eukaryotes, bacterial cells add another layer of complexity by concurrently replicating and segregating their chromosome (Toro et al., 2008; Marczynski et al., 2019). This concurrent replication and segregation require tight regulation to ensure each daughter cell receives intact copies of the genetic material. Chromosome replication is primarily regulated at the initiation step, which is driven by the canonical replication initiator DnaA (Messer, 2002). Various non-dedicated mechanisms have been proposed to drive the segregation of chromosome in bacteria (Gordon & Wright, 2000; Jun & Mulder, 2006; Jun & Wrigth, 2010; Yu et al., 2020), including DNA replication (Lemon & Grossman, 1998, 2000; Lemon et al., 2001), RNA transcription (Dworkin & Losick, 2002), and electrostatic repulsion of DNA (Brahmachari & Marko, 2019). Dedicated segregation machineries usually require nucleotide-dependent systems that are expected to work in conjunction with the entropic forces generated from non-dedicated mechanisms. Dedicated segregation mechanisms include the plasmid and chromosome partitioning system ParABS (Austin & Abeles, 1983; Ogura & Hiraga, 1983; Gerdes et al., 1985), nucleoid condensing proteins, such as SMC and MukB (Niki et al., 1991; Britton et al., 1998), and DNA translocases, such as FtsK and SpoIIIE, that clear DNA from the plane of division (Begg et al., 1995; Bath et al., 2000; Mahone et al., 2024).

The ParABS system is widely conserved (>70%) in bacteria (Livny et al., 2007). The system is composed of three components: 1) ParA, an ATPase protein that binds non-specifically to DNA (Havey et al., 2012; Vecchiarelli, Havey, et al., 2013; Vecchiarelli et al., 2014; Corrales-Guerrero et al., 2020); 2) ParB, a CTPase protein that binds DNA specifically at *parS* (Vecchiarelli & Funnell, 2013; Volante & Alonso, 2015; Tran et al., 2018; Osorio-Valeriano et al., 2019; Soh et al., 2019; Jalal & Le, 2020); 3) an origin-proximal chromosomal region, *parS*, where ParB binds (Livny et al., 2007; Toro et al., 2008; Havey et al., 2012). ParA binds non-specifically to DNA as an ATP-bound dimer that hydrolyzes ATP upon interactions with ParB-*parS* complexes, generating monomers that no longer bind DNA (Leonard et al., 2005; Vecchiarelli et al., 2010; Havey et al., 2012; Vecchiarelli, Havey, et al., 2013; Vecchiarelli, Hwang, et al., 2013; Hwang et al., 2013; Vecchiarelli et al., 2014; Volante & Alonso, 2015; Corrales-Guerrero et al., 2020). In addition to interactions with DNA and ParB, ParA has also been shown to interact with a variety of additional protein factors, many of which are localized at the cell poles. For example, in sporulating *Bacillus subtilis* cells, Soj (ParA) interacts with DivIVA, ComN, and RacA to tether the origins of replication to the membrane at the poles (Wu & Errington, 2003; Ben-Yehuda et al., 2003; Ben-Yehuda et al., 2005; dos Santos et al., 2012; Kloosterman et al., 2016). In *Streptomyces coelicolor,* ParA is recruited to the hyphal tips via Scy during the transition from hyphal cells to sporulating cells (Walshaw et al., 2010; Ditkowski et al., 2013). In *Vibrio cholerae*, HubP was shown to recruit both ParA and the origin to the cell poles (Yamaichi et al., 2012; Possoz et al., 2022). The existence of various protein partners found to directly interact with ParABS posit a multi-level system of regulation that is required to coordinate chromosome segregation with the progression of the cell cycle. However, our understanding of how ParA and its interactions with other proteins are coordinated over the cell cycle remains limited.

In *Caulobacter crescentus*, the ParABS system is essential for chromosome segregation (Mohl & Gober, 1997). The dimorphic life cycle of *C. crescentus* has allowed for extensive work on cell cycle regulation, including mechanistic details of the regulation of chromosome segregation and how cell polarity is determined and maintained (Figure 1A) (Collier, 2019; Govers & Jacobs-Wagner, 2020; Barrows & Goley, 2023). Prior to DNA replication initiation, the centromere-like region, *parS*, is anchored at the stalked (old) pole via ParB-PopZ and ParA interactions (Ebersbach et al., 2008; Bowman et al., 2008; Puentes-Rodriguez et al., 2023). Although DNA replication initiates at the origin of replication (*ori*), segregation does not initiate until *parS* (∼8 kb away from *ori*) is replicated (Toro et al., 2008; Hong & McAdams, 2011). A gradient-like structure of ParA extends from the new pole towards the stalk pole with the highest concentration of ParA at the new pole (Schofield et al., 2010; Ptacin et al., 2010). ParB promotes ParA’s ATPase activity that results in a retraction of the ParA gradient towards the new pole (Shebelut et al., 2010; Lim et al., 2014).

**Figure 1:**
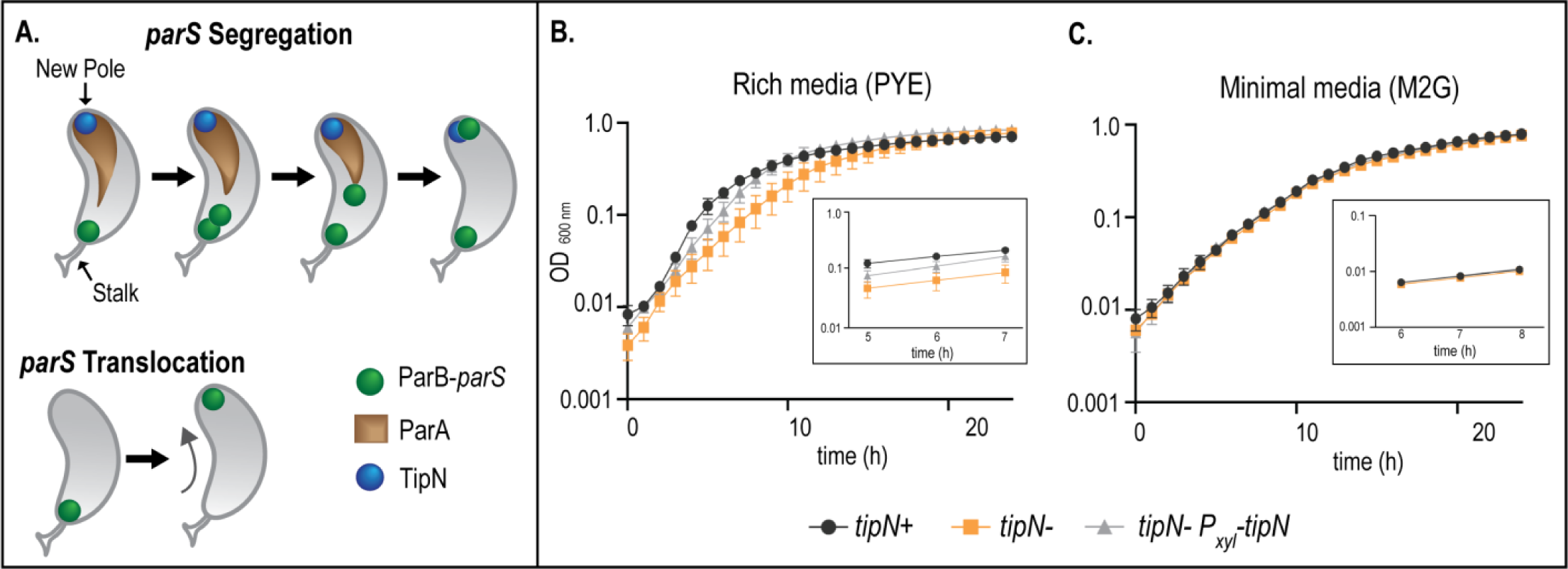
Loss of TipN has no impact on growth in minimal media. A) Schematic representation of *C. crescentus* highlighting ParB-*parS* (green), TipN (blue), and ParA dimers (brown) during chromosome replication and segregation of *parS*. (Bottom) Schematic of DNA-independent *parS* translocation. (B-C) Growth curves of wildtype (CB15N) cells expressing tipN (*tipN+*, black circles), Δ*tipN* (*tipN-*, orange squares), or with *tipN* complemented (*tipN-P_xyl_-tipN,* gray triangles) in (B) rich (PYE) media and in (C) minimal media supplemented with 0.1% xylose. Small insert represents exponential growth phase. Each graph is representative of three independent trials. Error bars are ± SD for three replicates per trial.

ParA has been shown to interact with two polar proteins in *C. crescentus*: TipN and PopZ (Schofield et al., 2010; Ptacin et al., 2010; Bowman et al., 2013; Ptacin et al., 2014; Holmes et al., 2016). The interactions between ParA and PopZ are critical for ParA’s ability to form a gradient and for PopZ to release the anchoring of *parS* from the cell pole for onset of segregation (Ptacin et al., 2014; Puentes-Rodriguez et al., 2023). TipN is a large, coiled-coil rich membrane-bound protein that localizes at the new pole during *parS* segregation (Lam et al., 2006; Huitema et al., 2006). TipN has been proposed to be involved in a variety of cell processes including the regulation of morphology, motility, and chromosome segregation (Lam et al., 2006; Huitema et al., 2006). The role of TipN-ParA interactions has remained debatable. For instance, TipN was shown *in vitro* to interact directly with ParA-ATP, favoring dimers versus monomers of ParA (Ptacin et al., 2010). However, *in vivo* analyzes of ParA variants in the presence and absence of TipN suggested TipN recruits monomers of ParA (Schofield et al., 2010). TipN is nonessential and *tipN* knockout cells display only subtle defects with chromosome segregation without severely impacting the ability of cells to grow and divide like wildtype cells (Lam et al., 2006; Schofield et al., 2010; Ptacin et al., 2010). Thus, TipN’s role with chromosome segregation remains unresolved.

In this study, we characterized the function of TipN in cells unable to replicate but capable of transporting un-replicated *parS* regions from the stalked pole to the new pole. Our data revealed that in the absence of chromosome replication, TipN becomes essential for ParA’s ability to transport *parS*. Using a *parA* merodiploid strain, we show that cells with increased levels of ParA can overcome the loss of *tipN*. Although TipN is non-essential, our work revealed that cells viability become susceptible to loss of TipN when the levels of the replication initiator DnaA are not regulated from the native promoter. Collectively, these findings confirm TipN’s role with chromosome segregation and highlight the complex regulation that is involved in the coordination of chromosome replication and segregation in bacteria.

## RESULTS

### ParA can translocate *parS* independent of DNA replication

One of the proposed functions of the polar protein TipN is with chromosome segregation and ParA. However, *tipN* knockout cells display subtle defects in chromosome segregation that mildly impact the cells’ ability to grow and divide (Lam et al., 2006; Ptacin et al., 2010). In fact, the subtle differences in doubling time between Δ*tipN* (99 ± 5 mins) and wildtype cells (93 ± 5 mins) are only observed in rich media (Figure 1B) (Lam et al., 2006). Our data show that Δ*tipN* cells have no defect in growth rates in minimal media compared to wildtype (Figure 1C). These data bring into question whether the polar factor TipN has a role with ParA’s ability to segregate *parS*. Our hypothesis was that chromosome replication contributes to ParA’s ability to segregate *parS,* obscuring the role of additional factors like TipN. If our premise were correct, removing those additional forces could reveal whether TipN is required for ParA to segregate *parS* from the stalked pole to the new pole. To test our hypotheses, we required conditions where ParA’s activity could be tested independent of chromosome replication. Thus, we used a strain where in the absence of chromosome replication, a sub-population of cells display migration of *parS* to the new pole (Mera et al., 2014). In this strain, chromosome replication is inhibited by depleting the replication initiator, DnaA, and the replication and segregation of *parS* is tracked with the fluorescently labeled ParB encoded at its native locus (*parB::cfp-parB P_van_-dnaA ΔvanA dnaA::Ω,* denoted here as *P_van_-dnaA*\*). To distinguish between the segregation of two replicated *parS* versus the transport of the single un-replicated *parS* from the stalked pole to the new pole, we will refer to the replication-independent transport of *parS* as translocation, as opposed to segregation (Figure 1A). In our analyzes of *parS* translocation in the absence of chromosome replication in the *P_van_-dnaA*\* strain, ∼ 30% of cells displayed *parS*-CFP-ParB localized at the new cell pole (Figure 2AE), consistent with previous reports (Melendez et al., 2019).

Because we are testing ParA’s activity and the potential connection with TipN, we examined whether ParA in the ParABS system is the sole mediator of *parS* transport when DNA replication is inhibited in the *P_van_-dnaA\** strain. We first tested the effect on *parS* translocation from increased levels of ParA by using a *parA* merodiploid with a second copy of *parA* under the xylose promoter. In the presence of xylose, this construct was shown to increase ParA levels ∼5x (Menikpurage et al., 2023). Cells expressing increased levels of wildtype ParA displayed an increase in the frequency of *parS* translocation from ∼25% in native ParA levels to ∼49% in cells with *parA* overexpressed (Figure 2DE). These data support the notion that the ParABS system is responsible for the translocation of *parS* in the absence of chromosome replication. To confirm that ParA was responsible for the observed replication-independent translocation of *parS*, we eliminated ParA’s activity by expressing a dominant negative *parA-K20R* variant that is incapable of promoting *parS* segregation (Toro et al., 2008). In the absence of inducer, both the empty vector control and cells encoding *P_xyl_-parA-K20R* retained similar frequency of *parS* translocation compared to cells expressing wildtype levels of ParA (Figure 2BCE). However, cells expressing *parA*-*K20R* display complete loss of *parS* translocation (1.6 ± 0.5%) (Figure 2CE). Collectively, these results demonstrate that the partitioning system ParABS retains its ability to transport *parS* to the new cell pole in the absence of DNA replication.

### ParA requires TipN to translocate *parS* in the absence of DNA replication

By removing chromosome replication as a contributing force to segregation, we then tested whether TipN is involved in ParA’s ability to segregate *parS*. To test the role of TipN with centromere segregation, we constructed a strain where the *tipN* gene was knocked out in the *P_van_-dnaA\** strain. We first confirmed that the *ΔtipN P_van_-dnaA*\* strain grown in the presence of the vanillate inducer displayed the previously reported chromosome segregation phenotype where *parS* is partially segregated (Ptacin et al., 2010). Consistent with the previous analyses, our data revealed a subpopulation of cells displaying partial *parS* segregation in the Δ*tipN* cells compared to *tipN*+ cells (Figure 3A). However, we observed a significantly lower percent of partial segregation (∼24%) in our analyses compared to what was previously reported (∼60%) (Ptacin et al., 2010). We posit that this difference is due to the genotype of the constructs used. The partial *parS* segregation analyzed previously was performed using a *parB* merodiploid strain (native *parB* and *P_van_-mCherry-parB*) to visualize the *parS*-ParB cellular localization (Ptacin et al., 2010), which results in higher ParB levels than wildtype. Altering the levels of ParB can impact its function with chromosome segregation (Mohl & Gober, 1997). The strain we used encodes the native *parB* fluorescently labeled at its native locus, which keeps the native levels of ParB undisturbed (Thanbichler & Shapiro, 2006). We can conclude that in the absence of TipN, a sub-population of cells display partial *parS* segregation, albeit not as severe phenotype as previously thought.

We then tested the effect of TipN on the ability of cells to translocate *parS* to the new pole in the absence of chromosome replication. Our data revealed that compared to the translocation observed in the presence of TipN, cells lacking *tipN* are unable to translocate *parS* (Figure 3B). The lack of *parS* translocation in Δ*tipN* cells resembled the lack of translocation in cells expressing inactive *parA-K20R* (Figure 2CE). To confirm that TipN was responsible for the loss of *parS* movement, we constructed a complementation strain with *tipN* under the xylose inducible promoter. Expression of *tipN* from the inducible promoter in Δ*tipN* cells rescues the ability to translocate *parS* to the opposite pole (Figure 3B). Collectively, these data support the involvement and potential requirement of TipN in *Caulobacter’s* ability to translocate *parS* to the opposite pole.

### High levels of ParA rescue *ΔtipN* segregation defects

To further analyze TipN’s requirement, we examined how ParA levels can impact *parS* segregation in cells lacking TipN. Because high levels of ParA increased the frequency of cells displaying *parS* translocation (Figure 2), we reasoned that increased levels of ParA could rescue the ability of cells to translocate *parS* in the absence of TipN. To test this hypothesis, we constructed a *tipN* knockout, *parA* merodiploid (native and *P_xyl_-parA)* in the *P_van_-dnaA\** background strain. Our data revealed that in a *tipN* knockout, cells grown in the presence of xylose to overexpress *parA* rescue the cell’s ability to translocate *parS* (Figure 3B). Cells that do not have an inducible copy of *parA* in *tipN* knockout cells were unable to translocate *parS* under the same conditions (Figure 3B). These data demonstrate that high levels of ParA can compensate for loss of TipN.

**Figure 2:**
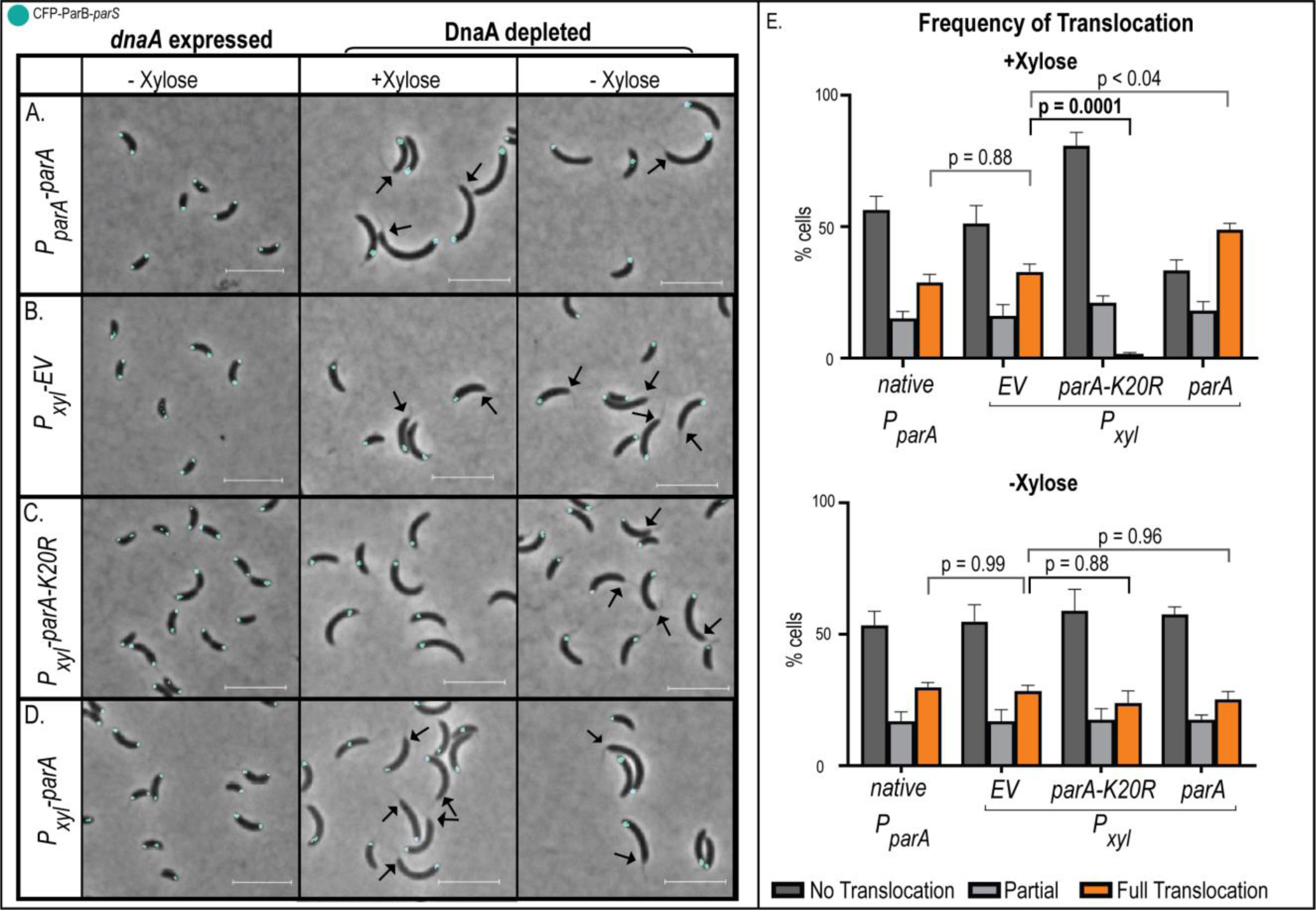
Increase levels of ParA results in increased replication-independent translocation frequency of *parS*. (A-D) Micrographs of mixed population of *C. crescentus* cells (*parB::cfp-parB, ΔvanA, P_van_-dnaA, dnaA::Ω*) expressing (A) *parA* from its native promoter (*P_parA_-parA)*, (B) *P_xyl_-empty vector (EV),* (C) *P_xyl_-parA-K20R,* and (D) *parA* overexpression (*P_xyl_-parA)* in minimal media. DnaA was either induced for expression or depleted for 3 hours prior to data analyses. Black arrows represent the location of the stalk in translocating cells. Scale bar – 5 µm. (E) Quantification of localization of the single CFP-ParB locus in A-D samples. Full translocation represents cells with CFP-ParB at the new cell pole. Data are mean averages from three independent trials. Error bars are ±SEM with two-way ANOVA analysis.

**Figure 3:**
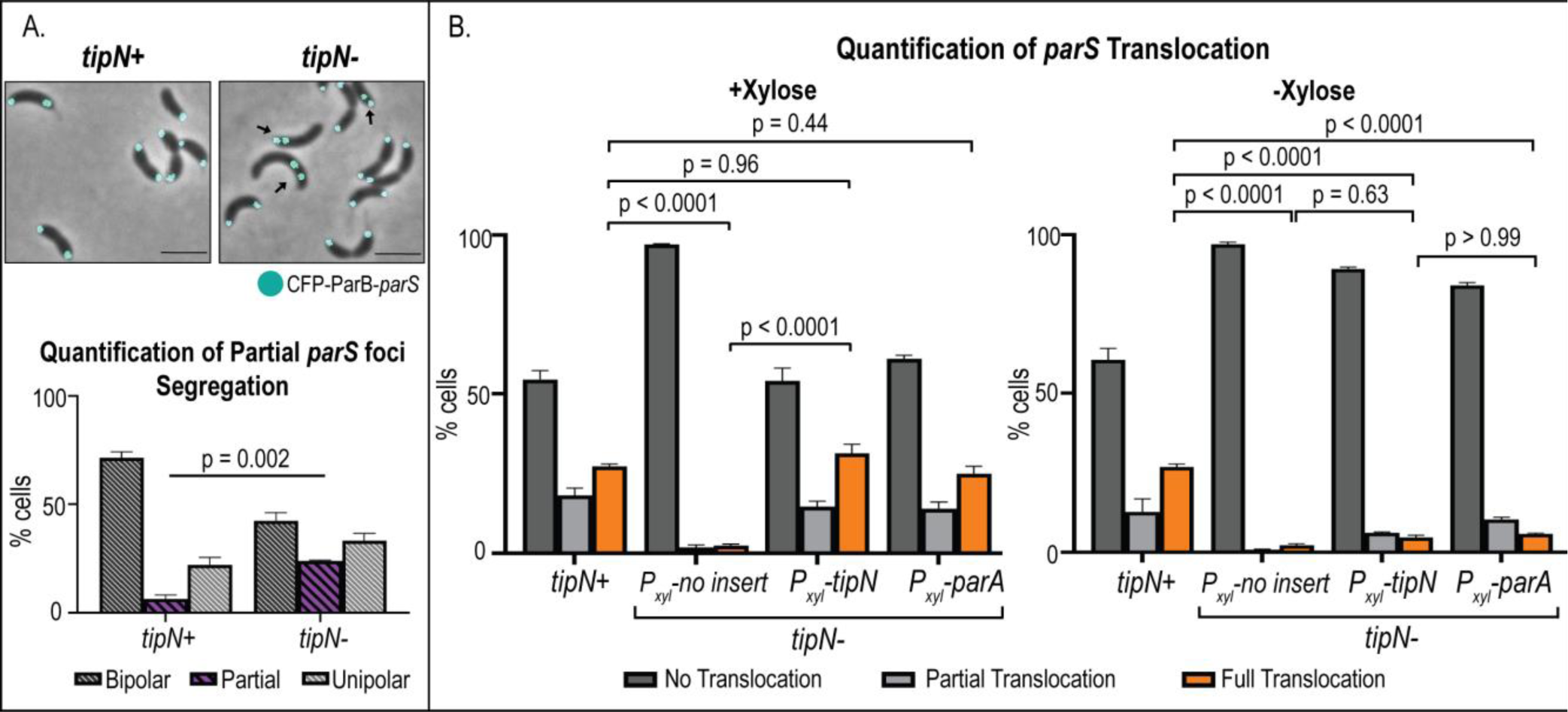
TipN is required for ParA to translocate *parS* in the absence of DNA replication. A) Loss of *tipN* cause a subtle but significant increase in cells with partial *parS* segregation. Micrographs of exponentially growing cells (*parB::cfp-parB, ΔvanA, P_van_-dnaA, dnaA::Ω* ± *tipN*) grown in minimal media supplemented with 100 μM vanillate. Scale bars – 2.5 µm. Black arrows indicate partial segregation. Quantification using blinded samples of the mean average percentage of cells with CFP-ParB at both poles (bipolar), one pole (unipolar), or in between (partial). B) Quantification of *parS* translocation after 3-hour DnaA depletion. Cells (*parB::cfp-parB, ΔvanA, P_van_-dnaA, dnaA::Ω*) included *tipN* expression from native promoter (*tipN*+), Δ*tipN* (*tipN*-), *tipN* expressed from the xylose promoter in Δ*tipN* (*tipN*-*P_xyl_*-*tipN*), or with overexpression of *parA* in Δ*tipN* (*tipN*-*P_xyl_*-*parA*). 0.1% xylose was added to induce *P_xyl_* expression. All data are from three independent trials. Error bars are ±SEM with 2-way ANOVA analyses.

### When DnaA levels are altered, cells become sensitive to loss of *tipN*

While analyzing TipN’s role with ParA, we identified a potential connection between TipN and the replication initiator, DnaA. In *C. crescentus*, the expression of *dnaA* is regulated over the cell cycle (Collier et al., 2006; Schrader et al., 2016; Frandi & Collier, 2019). However, *Caulobacter* cells engineered to express *dnaA* from inducible promoters have been widely used in the literature with no notable phenotypes. To confirm the lack of defects, we compared cell viability between wildtype (native expression) and cells expressing *dnaA* from the vanillate promoter. As expected, both strains display similar viability in our colony forming unit (CFU) analyses (Figure 4A). Similarly, TipN has been shown to be a non-essential protein for *Caulobacter*’s viability (Kirkpatrick & Viollier, 2014; Vellet et al., 2020). We confirmed that Δ*tipN* cells’ viability are indistinguishable compared to wildtype cells encoding *tipN* (Figure 4B). Interestingly, we found that Δ*tipN* cells expressing *dnaA* from an inducible promoter display a significant loss of viability compared to Δ*tipN* with *dnaA* expressed from its native promoter (Figure 4BC). This loss of viability required the combination of both Δ*tipN* and expression of *dnaA* from the inducible promoter (Figure 4C). We confirmed that the inducer (vanillate) itself was not impacting the viability of Δ*tipN* cells using CFU analyzes of cells grown in the presence of vanillate. Our data show that Δ*tipN* cells with native *dnaA* expression retain their viability just like wildtype cells when grown in the presence of vanillate (Figure 4B). To confirm that the viability defect of Δ*tipN* with *dnaA* expressed from an inducible promoter was due to the lack of TipN, we tested the complementation strain with *tipN* under the xylose inducible promoter. Cells expressing both *dnaA* and *tipN* from inducible promoters rescued the viability defect (Figure 4C). Collectively, these data revealed that altering the regulation of DnaA levels makes *C. crescentus* cells more susceptible to loss of TipN, suggesting a potential link between the regulation of replication initiation and centromere segregation.

**Figure 4:**
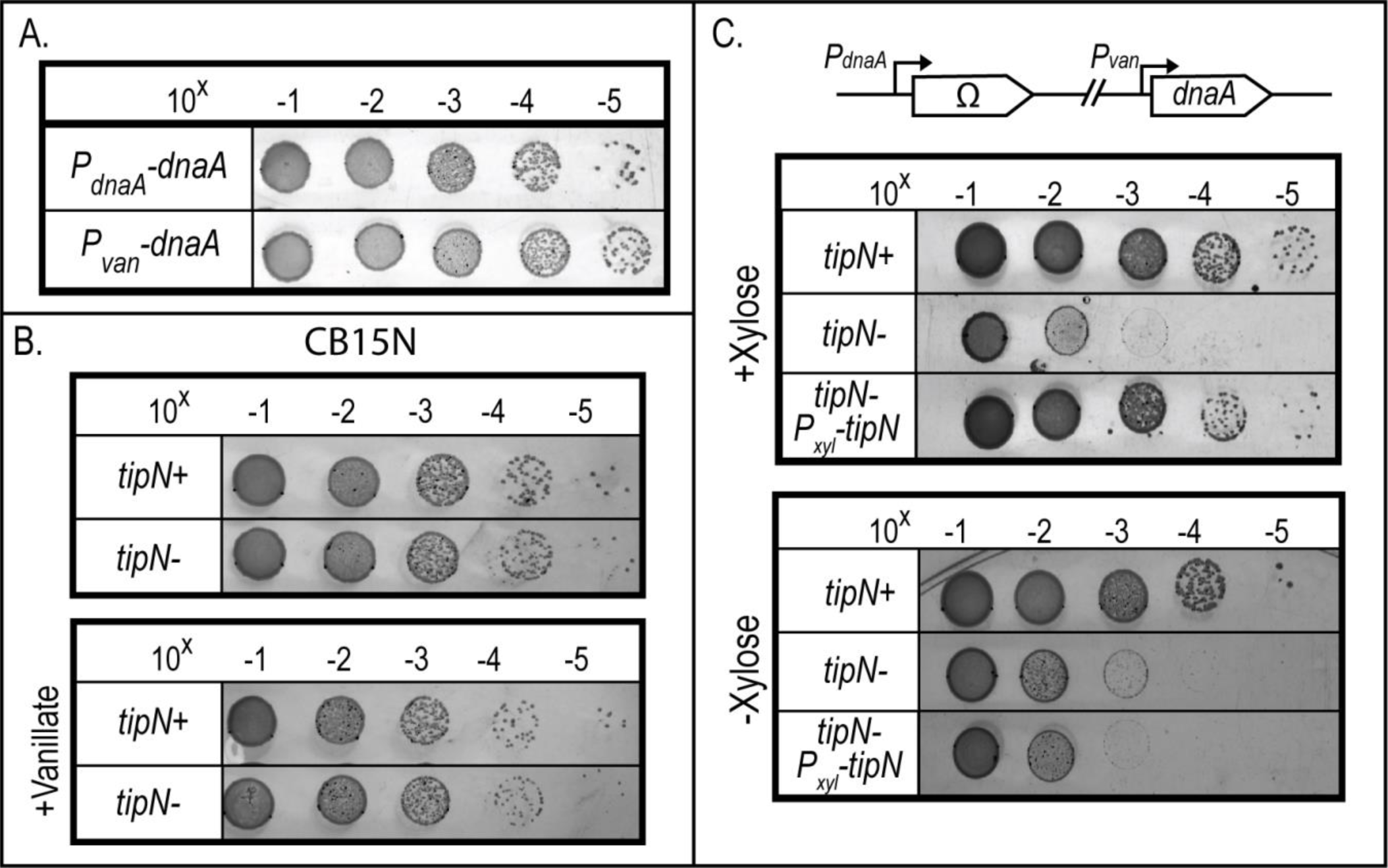
DnaA-dependent loss of viability of *ΔtipN* cells. Cell viability analyses using Colony forming unit (CFU) assays. Cells were serially diluted 1:10 and plated onto PYE plates supplemented with 100 µM vanillate, whenever noted. A) Comparison of cells expressing *dnaA* from native promoter (wildtype) and from inducible promoter vanillate (P_van_-dnaA) display no significant difference in CFUs. B) Comparison of wildtype cells CB15N cells with (*tipN*+) or without (*tipN*-) grown in the presence of vanillate reveals no impact on CFUs. C) The combination of *dnaA* expressed from inducible promoter (*P_van_*) and loss of *tipN* results in significant viability loss.

### The DnaA-dependent susceptibility of *ΔtipN* is likely connected to *parS* segregation

We examined whether the DnaA-dependent susceptibility of *ΔtipN* was due to problems with chromosome replication initiation and/or due to problems with centromere segregation. We first investigated potential problems with chromosome replication initiation because ParA in *C. crescentus,* as in other bacterial species, has been shown to impact the regulation of chromosome replication initiation (Murray & Errington, 2008; Kadoya et al., 2011; Menikpurage et al., 2023). Because *C. crescentus* initiates chromosome replication only once per cell cycle (Marczynski, 1999), we can track problems with chromosome replication initiation by simply quantifying the number of *oris* per cell. When the levels of ParA are increased, *C. crescentus* cells display more than 2 *oris* per cell (Menikpurage et al., 2023), albeit the mechanism of this connection remains unclear. We hypothesized that TipN could be the link that connects ParA to DnaA’s ability to initiate replication. To test our hypothesis, we examined the number of *oris* per cell in cells that overexpress *parA* but lack *tipN*. Based on the proximity between *parS* and *ori,* we used number of CFP-ParB foci per cell as proxy to represent number of *oris* per cell (Thanbichler & Shapiro, 2006; Toro et al., 2008). In cells expressing *dnaA* from the vanillate promoter, increased levels of ParA result in ∼ 30% of cells displaying >2 *oris* (Figure 5A). Under the same conditions, but using the Δ*tipN* background, the frequency of cells that display >2 *oris* remained the same as cells with *tipN+* (Figure 5A). These results revealed that TipN is not likely involved in the connection between ParA levels and regulation of chromosome replication initiation.

**Figure 5:**
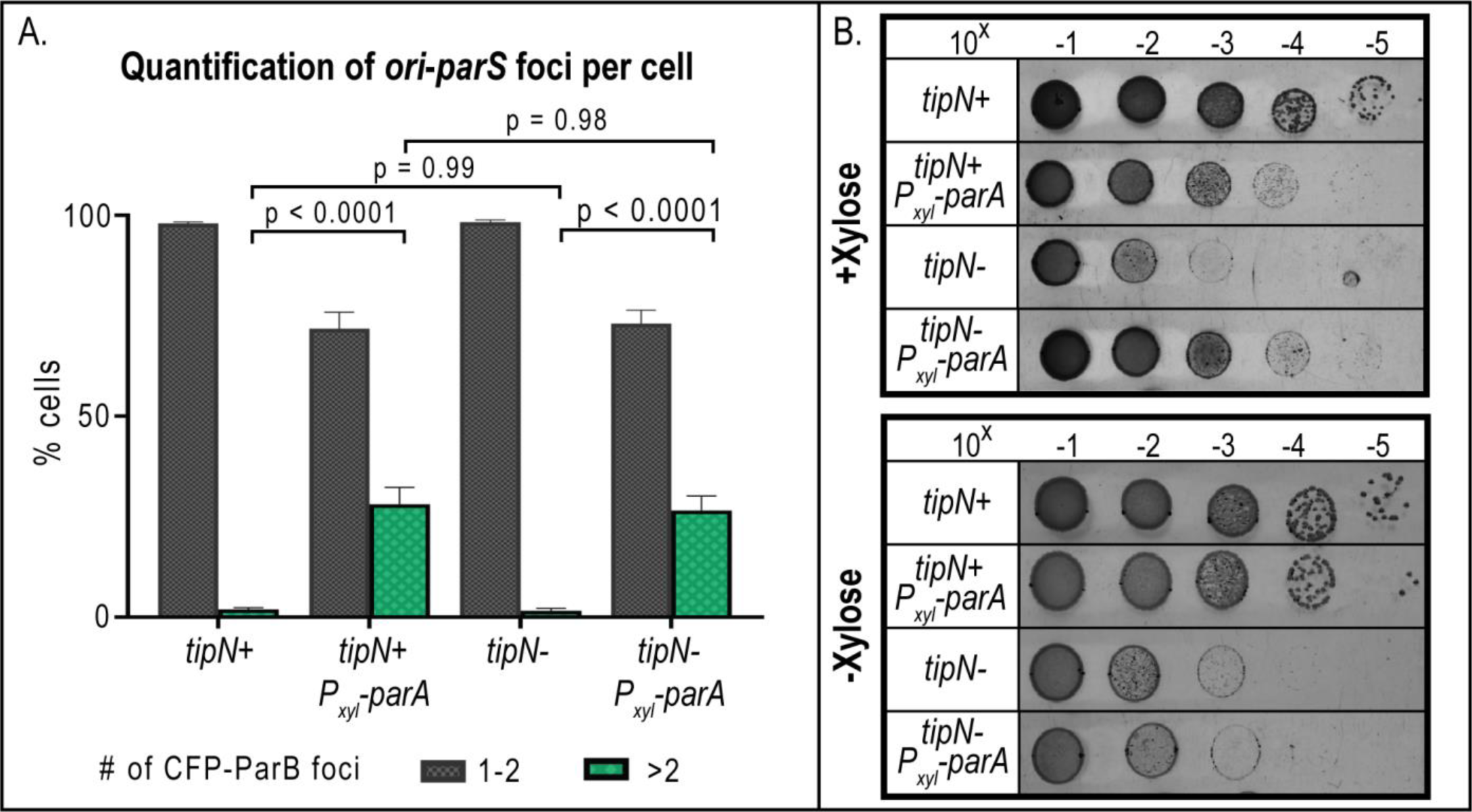
Characterization of DnaA-dependent susceptibility of *ΔtipN cells*. A) The increased number of cells with >2 *oris* is observed in cells with or without *tipN*. Quantification of the number of CFP-ParB foci to represent number of *ori* per cell. *P_xyl_-parA* was induced with 0.1% xylose for 3 hours in minimal media. Scale bars – 5 µM. Data are from three independent trials. Error bars are ±SEM with 2-way ANOVA analysis. B) Increased levels of ParA rescue DnaA-dependent viability defect of Δ*tipN* cells. Colony forming unit (CFU) viability assay in cells expressing *dnaA* from *P_van_* ± *tipN* and ± *P_xyl_-parA* induction. Plates were supplemented with 100 µM vanillate and 0.1% xylose was added to induce *P_xyl_* expression. CFUs are representatives of three independent trials.

We next examined whether the DnaA-dependent viability defect of Δ*tipN* cells was connected to problems with centromere segregation. We reasoned that if the problems were related to chromosome segregation, increasing levels of ParA could rescue the DnaA-dependent viability defect of Δ*tipN*. We based this reasoning on our results that increasing ParA levels rescue Δ*tipN* cells’ ability to translocate *parS* (Figure 3). To test our hypothesis, we first examined the impact on viability from increasing ParA levels in cells expressing *dnaA* from the vanillate promoter with *tipN* present at its native locus (Figure 5B). Consistent with the sub-population of cells that over-initiate chromosome replication (Figure 5A), we observed a loss of viability that can be attributed to the high levels of ParA and over-initiation of replication. For the Δ*tipN* cells, increasing the levels of ParA rescue the DnaA-dependent viability defect (Figure 5B). Collectively, these data suggest that the DnaA-dependent viability defect of Δ*tipN* cells is likely due to problems related with *parS* segregation. This potential new connection between TipN, ParA, and DnaA requires further investigation to identify mechanistic details of the complex regulation that drives the coordination between onset of chromosome replication and segregation.

## DISCUSSION

The ability of bacteria to maintain intact copies of their chromosome is crucial for their survival. Most bacteria use the partitioning ParABS system to direct chromosome segregation. Proteins that localize at the cell poles have also been implicated with chromosome segregation, in many cases directly interacting with the ParABS system. In *Caulobacter crescentus*, the polar factor TipN interacts with ParA and has been proposed to be involved with chromosome segregation. By inhibiting chromosome replication, a non-ParABS contributor of chromosome segregation, our analyses revealed that TipN is required for ParA’s ability to segregate the centromeric region *parS*. TipN’s requirement for *parS* segregation can be compensated by increasing the cellular levels of ParA. We identified a DnaA-dependent susceptibility of Δ*tipN* that is likely due to problems with *parS* segregation. Our work provides new insights into the combined roles between DNA replication and the partitioning system ParABS in the transport of the *ori*-proximal chromosomal region, *parS*.

Our data revealed the significant role that the entropic forces from chromosome replication or the process of chromosome replication itself (Lemon et al., 2001; Dworkin & Losick, 2002; Jun & Mulder, 2006; Jun & Wrigth, 2010) play with the partitioning system’s ability to transport *parS*. In the presence of chromosome replication, the role of the polar protein TipN with *parS* segregation is relatively disposable. This is evident in the ability of cells without *tipN* to grow in minimal media at the same doubling rates as cells with *tipN* (Figure 1C). However, in the absence of chromosome replication this disposable function of TipN becomes essential. ParA was unable to transport *parS* in *tipN* knocked out cells in the absence of chromosome replication (Figure 3B). Thus, we envision a model where TipN in normally replicating cells plays a fine-tuning role, likely in ParA’s directionality of transport, as previously proposed (Schofield et al., 2010; Ptacin et al., 2010). This potential fine-tuning role would become more important when cells are growing and coordinating cell cycle progression at faster rates, as is the case when cells are grown in rich media. This would be consistent with the growth defect in Δ*tipN* cells grown in rich media (Lam et al., 2006) (Figure 1B).

The observation that cells expressing the replication initiator *dnaA* from a non-native promoter lose viability in the absence of *tipN* suggest that TipN may also be involved in the complex coordination between replication initiation and centromere segregation. The many examples that have been reported about the connections between replication initiation and chromosome segregation highlight the significance of maintaining these two critical events highly coordinated. In *Caulobacter*, replication was shown connected to segregation via the nucleoid-associated protein, GapR (Taylor et al., 2017). Cells depleted of *gapR* experienced delayed chromosome replication initiation and mis-localized the centromere, *parS* (Taylor et al., 2017). Furthermore, changing the levels of ParA or ParB alters the numbers of origins of replication per cell (Mohl & Gober, 1997; Mohl et al., 2008; Menikpurage et al., 2023). DnaA itself can disrupt the onset of centromere segregation by directly binding at *parS* (Mera et al., 2014). The connection between replication and segregation has also been shown in other bacterial species. For instance, Bacillus ParA (Soj) can activate or directly deactivate the activity of DnaA (Murray & Errington, 2008; Kadoya et al., 2011; Scholefield et al., 2012). The tight coordination between replication initiation and chromosome segregation is likely widely conserved given that the *parS* chromosomal locus is usually found near the origin of replication in the diverse bacterial species that encode the partitioning system ParABS (Livny et al., 2007).

Our data confirmed that TipN is involved in chromosomal maintenance during the progression of the cell cycle. Notably, polar proteins that have been shown to associate with the ParABS system commonly have coiled-coil domains. In *C. crescentus*, the cytoplasmic domain of TipN, which includes the coiled-coil domain, was shown to directly interact with ParA (Ptacin et al., 2010). In *Streptomyces*, the polar protein Scy was shown to interact with ParA through its coiled-coil domains and this interaction has been proposed to regulate ParA’s oligomerization (Walshaw et al., 2010; Ditkowski et al., 2013). In *Bacillus* and mycobacteria, DivIVA interacts with ParA/Soj via the coiled-coil domain (Ginda et al., 2013; Pioro et al., 2019). In *Vibrio*, *Shewanella*, *Photobacterium*, and several other gammaproteobacteria, the polar protein HubP recruits ParA-type protein partners whose functions are involved with chromosome segregation, motility, and chemotaxis systems (Yamaichi et al., 2012; Takekawa et al., 2016; Possoz et al., 2022; Altinoglu et al., 2022). Using coiled-coil structural domains for regulating chromosome segregation is not restricted to bacteria. The canonical structural maintenance of chromosomes (SMC) family of proteins conserved in all domains of life include coiled-coil domains and are central regulators of chromosome segregation (Hirano, 2005; Yanagida, 2005; Uhlmann, 2016; Truebestein & Leonard, 2016; Yatskevich et al., 2019). Coiled-coil proteins are also found in protein structures involved in chromosome segregation in eukaryotes, like in kinetochores (Truebestein & Leonard, 2016). In sum, the use of coiled-coil domains in the regulation of chromosomal maintenance is widely conserved.

## MATERIALS AND METHODS

### Strains and plasmids

All the strains used in this study are listed in Tables 1. Plasmids were constructed by cloning the PCR amplified DNA fragments using genomic DNA from wild-type *C. crescentus* CB15N (NA1000) as the PCR template. PCR amplicons were then placed into pNTPS138 or pXCHYC-2 vectors (Thanbichler et al., 2007) via Gibson assembly (Gibson, 2009; Gibson et al., 2009) or restriction enzyme cloning and ligation. Plasmids were transformed via heat shock into *E. coli* DH5α cells and grown in Luria-Bertani (LB) medium at 37°C with shaking. Plasmid constructs were verified via sequencing. Mobilization of the plasmid into *C*. *crescentus* was done via electroporation.

**Table 1.** *Caulobacter crescentus* strains used in this study.

| Strain ID | Genotype | Reference |
| --- | --- | --- |
| CB15N | Synchronizable derivative of *C. crescentus* CB15 (NA1000) | Evinger, M. and N. Agabian, 1997 |
| PM109 | CB15N, *parB::cfp-parBΔvanA P<sub>van</sub>-dnaA-aaCl dnaA::Ω (spec<sup>R</sup>/strep<sup>R</sup>)* | Mera et al., 2014 |
| PM542 | CB15N, *parB::cfp-parBΔvanA P<sub>van</sub>-dnaA-aaCl dnaA::Ω (spec<sup>R</sup>/strep<sup>R</sup>) P<sub>xyI</sub>-parA<sup>WT</sup>* | Menikpurage et al., 2023 |
| PM747 | CB15N, *parB::cfp-parBΔvanA P<sub>van</sub>-dnaA-aaCl dnaA::Ω (spec<sup>R</sup>/strep<sup>R</sup>) ΔtipN* | This study |
| PM771 | CB15N, *parB::cfp-parBΔvanA P<sub>van</sub>-dnaA-aaCl dnaA::Ω (spec<sup>R</sup>/strep<sup>R</sup>) P<sub>xyI</sub>-empty vector (EV)* | Menikpurage et al., 2023 |
| PM918 | CB15N, *parB::cfp-parBΔvanA P<sub>van</sub>-dnaA-aaCl dnaA::Ω (spec<sup>R</sup>/strep<sup>R</sup>) P<sub>xyI</sub>-par-K20R* | Menikpurage et al., 2023 |
| LS4527 | CB15N, *ΔtipN* | Lam et al., 2006 |
| PM979 | CB15N, *parB::cfp-parBΔvanA P<sub>van</sub>-dnaA-aaCl dnaA::Ω (spec<sup>R</sup>/strep<sup>R</sup>) P<sub>xyI</sub>-parA<sup>WT</sup>* | This study |
| PM1096 | CB15N, *parB::cfp-parBΔvanA P<sub>van</sub>-dnaA-aaCl dnaA::Ω (spec<sup>R</sup>/strep<sup>R</sup>) ΔtipN P<sub>xyI</sub>-tipN* | This study |

### Growth Conditions and Analyses

*C. crescentus* strains were inoculated from freezer stocks and grown in minimal media (M2G) or rich media (PYE) at 30°C, as previously described (Ely, 1991). Exponentially growing cultures were used for all experiments. For *C. crescentus,* liquid media was supplemented with antibiotics at the following concentrations: 5 μg/mL kanamycin, 25 μg/mL spectinomycin, 5 μg/mL streptomycin, or 1 μg/mL gentamycin. PYE plates were supplemented with 25 μg/mL kanamycin, 100 μg/mL spectinomycin, 5 μg/mL streptomycin, or 5 μg/mL gentamycin. Where necessary, 100 μM vanillic acid and/or 0.1% xylose was added to induce expression from the vanillate or xylose promoters (*P_van_* or *P_xyl_*), respectively (Thanbichler et al., 2007). *E. coli* strains were inoculated from freezer stocks and grown in LB media at 37°C. For *E. coli*, liquid media was supplemented with 30 μg/mL kanamycin. LB plates were supplemented with 50 μg/mL kanamycin. *Growth analyses.* Early log phase cultures of *C. crescentus* strains were used to inoculate in minimal media (M2G) or rich media (PYE) in 96-well plates. Absorbances at OD_600_ were measured every hour for 24 hours in a Biotek EPOCH-2 microplate reader at 30°C with orbital shaking at 180 rpm.

### DnaA Depletion analyses

*C. crescentus* strains were inoculated from freezer stocks into 2 mL of M2G media and grown to mid-log phase (OD_600_ ∼ 0.4-0.5) and then re-inoculated into 10 mL of M2G media and grown overnight at 30°C to early log phase (OD_600_ ∼ 0.3). Media was supplemented with 100 μM vanillic acid to induce production of DnaA (*P_van_-dnaA*) (Thanbichler et al., 2007). Cultures were washed three times in 1 mL of 1x M2 salts to remove inducer. Following the washes, cells were resuspended in 2 mL of minimal (M2G) media ± 100 µM vanillic acid to induce *P_van_* and ± 0.1% xylose to induce *P_xyl_*. Cells were spotted onto 1% agarose pads for analyses. The location or number of CFP-ParB was manually counted using the “Cell Counter” plugin from ImageJ/FIJI (Schindelin et al., 2012). These assays were done with blinded samples and in triplicate with at least 100 cells quantified per condition per replicate. The statistical analysis performed was 2-way ANOVA with multiple comparisons.

### Plating Colony Forming Units for Viability

*C. crescentus* strains were inoculated from freezer stocks and grown in minimal media (M2G) or rich media (PYE) at 30°C and grown overnight to early log phase (OD_600_<0.3) and normalized to OD_600_ = 0.1. Each culture was then serially diluted 1:10 in fresh media in a 96-well plate. 5 μL of each dilution was spotted onto PYE plates supplemented with 100 µM vanillic acid and/or 0.1% xylose, where appropriate. Plates were grown at 30°C for two days and then imaged.

### Fluorescence Microscopy

Live cells (∼2 μL) were immobilized on M2G + 1% agarose pads. Live cells were imaged at room temperature (RT, ∼22°C). Phase contrast and epifluorescence images were obtained using the Zeiss Axio Observer 2.1 inverted microscope with AxioCam 506 mono camera (objective: Plan-Apochromat 100x/1.40 oil Ph4 M27 [WD=0.17mm]) and Zen Pro software. CFP-ParB was imaged using the CFP filter.

